# *Ralstonia solanacearum* alters root developmental programs in auxin-dependent and independent manners

**DOI:** 10.1101/2022.06.22.497157

**Authors:** Lu Zhang, Gang Yu, Hao Xue, Meng Li, Rosa Lozano-Durán, Alberto P. Macho

## Abstract

Microbial pathogens and other parasites can modify the development of their hosts, either as a target or a side effect of their virulence activities. The plant pathogenic bacterium *Ralstonia solanacearum*, causal agent of the devastating bacterial wilt disease, is a soil-borne microbe that invades host plants through their roots, and later proliferates in xylem vessels. In this work, we studied the early stages of *R. solanacearum* infection in the model plant *Arabidopsis thaliana*, using an *in vitro* infection system. In addition to the previously reported inhibition of primary root length and increase in root hair formation at the root tip, we observed an earlier xylem differentiation during *R. solanacearum* infection that occurs in a HrpG-dependent manner, suggesting that the pathogen actively promotes the development of the vascular system upon invasion of the root. Moreover, we found that the phytohormone auxin, of which the accumulation is promoted by the bacterial infection, is required for the *R. solanacearum*-triggered induction of root hair formation, but not earlier xylem differentiation. Altogether, our results shed light on the capacity of *R. solanacearum* to induce alterations of root developmental pathways and on the role of auxin in this process.

## INTRODUCTION

Upon invasion of a host plant, microbial pathogens can modify plant development in order to facilitate plant colonization and/or pathogen proliferation. *Ralstonia solanacearum* is a pathogenic bacterium that causes bacterial wilt disease in more than 250 plant species, including agriculturally important crops such as tomato, potato, banana, and peanut (Genin, 2010; Mansfield *et al*., 2012). *R. solanacearum* is soil-borne, and can invade plants through the roots, using wounds, root tips, and secondary root emerging points as penetration sites. Upon plant invasion, bacteria move through the root cortex until they reach the vascular system, where they use the xylem vessels to proliferate and spread systemically in the plant (Xue *et al*., 2020). The pathogenicity of *R. solanacearum* relies on several virulence factors, including a type III secretion system (T3SS), which secretes effector proteins (type III effectors; T3Es) inside plant cells in order to manipulate plant cellular functions and suppress plant immunity (Boucher *et al*., 1985; Macho, 2016). In *R. solanacearum*, the expression of the components of the T3SS and of the T3Es is regulated by HrpB, while the central virulence regulator HrpG controls the expression of HrpB and other pathogenicity determinants (Valls *et al*., 2006).

Plant roots are essential organs and mediate the interaction with soil-borne bacteria. In the model plant *Arabidopsis thaliana* (hereafter, Arabidopsis), the root system can be divided in primary root, which is formed embryonically, and secondary roots, which develop post-embryonically (Scheres *et al*., 1994). The primary root can be further divided into four developmental zones: meristematic, transition, elongation, and maturation zones, based on their cellular activities (Motte *et al*., 2019). The root meristem contains the stem cell niche that gives rise to different cell types and the generated daughter cells. There are different types of stem cells in the stem cell niche: columella stem cells, lateral root cap/epidermis initials, cortex/endodermal initials, and vascular initials (Dolan *et al*., 1993). A small group of mitotically less active cells, named quiescent center (QC), is located at the center of the stem cell niche (Motte *et al*., 2019), and plays a key role in its positioning and maintenance. Primary root development is controlled in the stem cell niche, which is established during embryogenesis (Scheres *et al*., 1994). Upon leaving the meristem, cells modify their physiological state to initiate a rapid elongation. However, in the transition zone, which is located shootward of the meristematic zone, the cells do not divide, and cell lengthening is relatively slow compared with that in the neighboring elongation zone (Verbelen *et al*., 2006). In the elongation zone, cells reach a period of elongation and experience a series of cellular alterations. After cells extend to their final size, they enter the maturation zone and acquire their mature and unique characteristics. There are two distinguishable features of the maturation zone. The first one is the formation of the Casparian strip, which functions as a protective barrier to prevent apoplastic entry of water and solutes into the central vascular cylinder (Roppolo *et al*., 2011), and xylem. The other notable feature of the maturation zone is the appearance of root hairs, which emerge from epidermal cells overlying two cortical cells (Duckett *et al*., 1994).

*In vitro* inoculation systems in model plants, such as Arabidopsis, *Medicago truncatula*, petunia, and tomato, have allowed the characterization of macroscopic alterations in root development caused by *R. solanacearum* infection. Among them, the most striking alteration consists in a strong inhibition of primary root growth, observed in several plant species, and likely caused by cell death at the root tip and dysfunction of the root apical meristem (Zolobowska & Van Gijsegem, 2006; Turner *et al*., 2009; Digonnet *et al*., 2012; Lu *et al*., 2018). Moreover, *R. solanacearum* infection induces the formation of root hairs at the root tip (Lu *et al*., 2018). Although the formation of lateral roots seems to be inhibited at early stages of *R. solanacearum* infection (Zolobowska & Van Gijsegem, 2006; Digonnet *et al*., 2012), it is enhanced at late stages (Zhao *et al*., 2019); it could be hypothesized that the early inhibition of lateral roots may be a consequence of early defence responses, while bacterial manipulation may promote the formation of lateral roots at later stages, following bacterial proliferation. Interestingly, the formation of root hairs and lateral roots may provide additional penetration sites for *R. solanacearum* cells in the rhizosphere, hence potentially benefiting the bacterium at the population scale. At the microscopic level, *R. solanacearum* infection seems to promote the formation of mature xylem vessels close to the root tip upon inoculation in *M. truncatula* (Turner *et al*., 2009). Whether these developmental alterations are the result of direct virulence activities aimed at promoting bacterial pathogenesis or are just a collateral effect of the events enabling or derived from the microbial invasion remains to be determined; the nature of these changes, however, according to the current understanding of the bacterial invasion process, supports the former. Altogether, it could be hypothesized that *R. solanacearum* alters root development to facilitate root invasion or plant colonization, although the molecular mechanisms underpinning these alterations remain unknown.

The phytohormone auxin is a plant growth regulator essential for the development of the root system (Benková & Hejátko, 2008; Overvoorde *et al*., 2010; Motte *et al*., 2019). The establishment of auxin gradients in the root requires the interplay of local auxin biosynthesis (Brumos *et al*., 2018), transport, and signaling. Auxin is supposed to accumulate mainly in the primary root tip by polar transport from the shoot, which mainly depends on the action of directional transporters such as the PINs (Gälweiler *et al*., 1998; Blilou *et al*., 2005; Grieneisen *et al*., 2007). In Arabidopsis, tomato, and wild potato, the expression of auxin-related genes is upregulated upon *R. solanacearum* inoculation (Zuluaga *et al*., 2015; French *et al*., 2018; Zhao *et al*., 2019). Interestingly, a tomato resistant cultivar displayed an exclusive downregulation (not observed in a susceptible cultivar) of auxin-related genes during *R. solanacearum* infection (French *et al*., 2018), which supports a potential association of auxin responses with bacterial virulence or plant resistance; however, auxin-related genes were upregulated in both resistant and susceptible wild potato cultivars upon inoculation with *R. solanacearum* (Zuluaga *et al*., 2015). Supporting the role of auxin for the *R. solanacearum*-induced developmental alterations, it has been shown that *R. solanacearum* does not induce the formation of root hairs in the Arabidopsis auxin-insensitive *tir1* mutant, although this mutant still shows inhibition of root growth and cell death at the root tip after bacterial inoculation (Lu *et al*., 2018). Moreover, a tomato *dgt1-1* mutant, which has altered auxin transport and impaired formation of secondary roots, exhibits enhanced resistance to *R. solanacearum* (French *et al*., 2018). Therefore, it is tempting to hypothesize that auxin-mediated morphological alterations actively contribute to the success of *R. solanacearum* infection.

Here, we sought out to identify the developmental alterations of the Arabidopsis root caused by *R. solanacearum* infection at the microscopic level and the signaling pathways underlying the observed changes. For this purpose, we combined confocal imaging, fluorescent gene expression markers, and pharmacological and genetic approaches. Our results indicate that invasion by this bacterial pathogen alters root developmental programs and modifies root architecture in a HrpG-dependent manner, and that some of these changes (root hair formation), but not all (premature xylem differentiation), depend on the impact of *R. solanacearum* on auxin accumulation.

## RESULTS

### *R. solanacearum* infection induces root hair formation and xylem differentiation

In order to dissect different macroscopic and microscopic alterations in Arabidopsis roots in early stages after *R. solanacearum* inoculation, we used an *in vitro* inoculation assay, similar to that described by Lu *et al* (Lu *et al*., 2018). In this assay, we inoculated Arabidopsis seedlings using a low inoculum dose (10^5^ cfu/ml, compared to 10^7^ cfu/ml used in (Lu *et al*., 2018)) by placing a 5 microliters droplet of bacterial inoculum 1 cm above the root tip. Similar to what was previously described (Lu *et al*., 2018), inoculation with the *R. solanacearum* wild-type reference strain GMI1000 caused an immediate reduction of root growth (Figure 1A and 1B) and an increase in the formation of root hairs (Figure 1C and 1D) in comparison with mock-treated roots. On the contrary, an *hrpG* mutant strain, lacking a major regulator of the expression of several virulence factors, including the T3SS (Valls *et al*., 2006), did not cause any significant alterations in Arabidopsis roots (Figure 1A-D). To visualize alterations at the microscopic level, we stained roots with propidium iodide (PI), a fluorescent dye that stains plant cell walls, and observed root tips using confocal microscopy. Interestingly, we observed an earlier differentiation of xylem cells in the elongation zone of roots inoculated with GMI1000, compared to mock- or *hrpG*-inoculated roots (Figure 1E), measured by a significantly shorter distance between the QC and the first appearance of differentiated xylem (Figure 1F), suggesting that *R. solanacearum* induces earlier xylem differentiation in a T3SS-dependent manner. However, given that an active *R. solanacearum* infection causes a reduction in root growth, we considered the possibility that the proximity between the QC and the first appearance of differentiated xylem is simply a consequence of reduced meristem size; nevertheless, the observation that the number of meristematic cortex cells did not significantly vary between seedlings inoculated with water, GMI1000, or *hrpG* (Figure 1G and 1H), suggests that this is not the case. We also considered the possibility that differences in cell elongation before differentiation could account for both the inhibition of root growth and the earlier xylem differentiation. However, our results show an apparent reduction in the length of the second and third cortex cells in the elongation zone in both GMI1000- and *hrpG*-inoculated seedlings (Figure 1I), which indicates a general impact of bacterial presence in the root, and suggests that differences in cell elongation do not underlie the differences in cell differentiation caused by GMI1000 inoculation. The formation of the Casparian strip in the endodermis is another hallmark of tissue differentiation in roots. Seedlings inoculated with GMI1000 show an apparent reduction in the distance from the QC to the appearance of the Casparian strip (Figure 1J and 1K), which indeed suggests an earlier differentiation of the endodermis. However, when we visualize the Casparian strip and xylem lignification simultaneously, it becomes evident that, while these structures appear concomitantly in untreated roots, their formation is uncoupled in inoculated roots (Figure S1I and S1J), indicating a specific effect on xylem differentiation that goes beyond a potential indirect effect caused by the inhibition of root growth.

**Figure 1.**
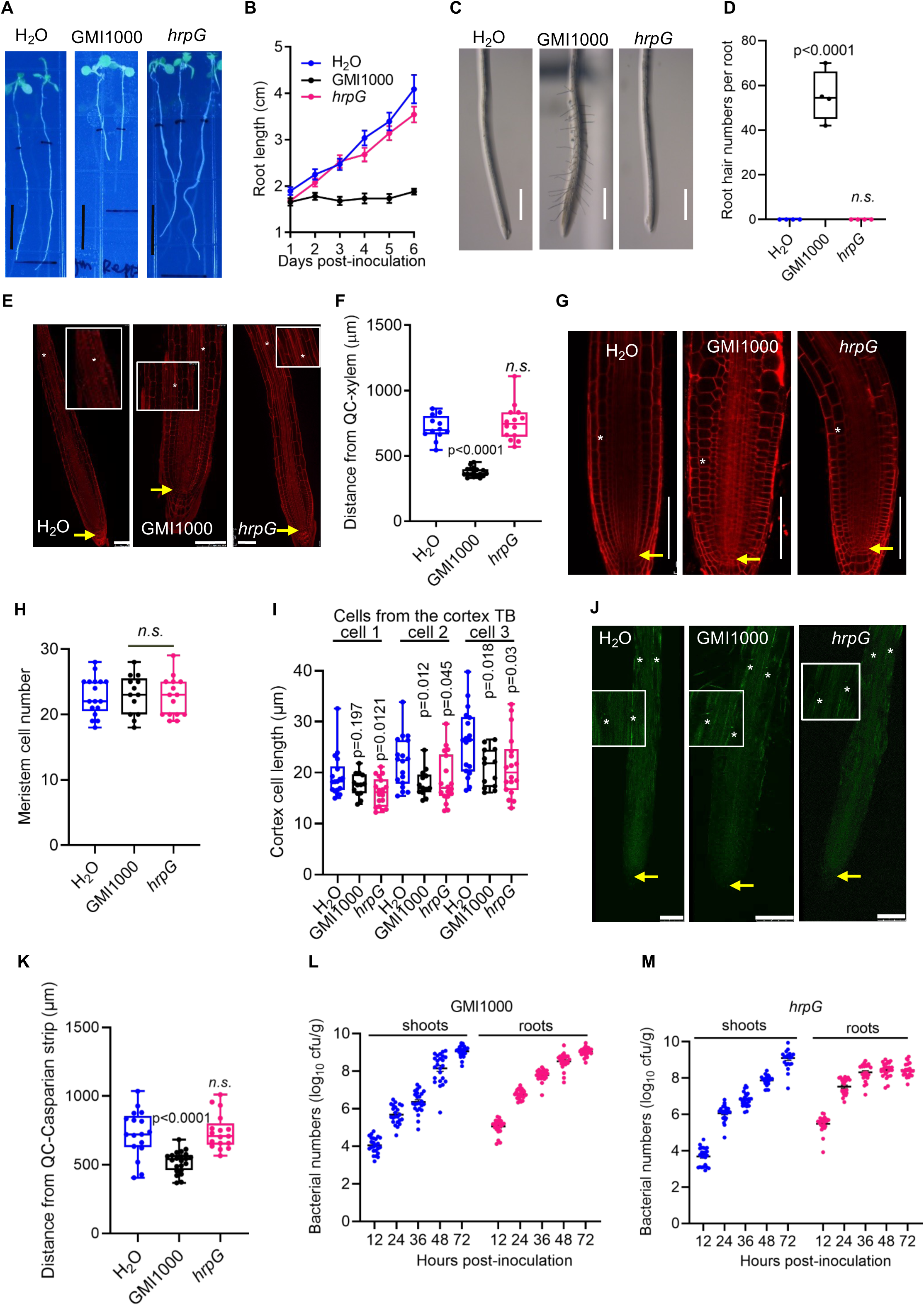
Inoculation with *Ralstonia solanacearum* causes Arabidopsis root development alterations. (A) *R. solanacearum* GMI1000 inhibits primary root growth in Arabidopsis. The pictures were taken 6 days post-inoculation (dpi), scale bar=1 cm. (B) Time-course observations of primary root growth in different days post-inoculation. Data are represented as means ± SEM (n=8 in each condition). (C) *R. solanacearum* GMI1000 induces root hair formation in Arabidopsis. Pictures were taken 54 hours post-inoculation (hpi), scale bar= 500 μm. (D) Quantification of root hairs in (C) (n=4 in each condition). Root hairs were counted in the 1.5-2 mm interval from the root tip in each root. (E and F) *R. solanacearum* GMI1000 causes earlier xylem differentiation in Arabidopsis. In (E), the inoculated seedlings were stained with propidium iodide (PI) at 54 hpi. Yellow arrows and white asterisks indicate the quiescent center (QC) and the first appearance of xylem, respectively, scale bar=100 μm. In (F), the distance from the QC to the xylem was measured from (E), and data are represented as pools of three repeats. (G and H) *R. solanacearum* GMI1000 does not affect root meristem cell numbers in Arabidopsis. In (G), the inoculated seedlings were used for PI staining at 54 hpi. Yellow arrows and white asterisks indicate the QC and the first cortical cell that was twice the length of the preceding cell, respectively, scale bar=100 μm. In (H), meristem cells were counted and data are represented as pools of three repeats. (I) Measurement of cell length of newly differentiated cortex cells in (G). TB means cortex transition boundary (TB), and the measurements start from the first cell indicated with white asterisk. Data are represented as pools of three repeats. (J and K) *R. solanacearum* GMI1000 causes earlier casparian strip formation in Arabidopsis. In (J), the inoculated seedlings were de-stained at 54 hpi and the auto-fluorescence of dot-like appearance of Casparian strip formation were observed. Yellow arrows and white asterisks indicate the quiescent center (QC) and the first appearance of Casparian strip, respectively, scale bar=100 μm. In (K), the distance from the QC to the first appearance of Casparian strip was measured from (J), and data are represented as pools of three repeats. (L) Quantification of *R. solanacearum* GMI1000 growth in different tissues in Col-0 seedlings. (M) Quantification of *R. solanacearum hrpG* mutant growth in different tissues in Col-0 seedlings. In all experiments, 7-d-old Arabidopsis Col-0 seedlings grown on MS plates were inoculated with different *R. solanacearum* strains using a 1×10 cfu/ml inoculum. In all figure panels, GMI1000 means *R. solanacearum* GMI1000 wild-type strain; *hrpG* means *R. solanacearum* GMI1000 *hrpG* mutant strain; H_2_O means mock (in which water has been used instead of the bacterial inoculum). Each experiment was repeated at least three times, except for (C and D, two times) with similar results. In (D, F, H, I, K-M), data are represented as boxes and whiskers, center line, median; box limits, lowest and highest data point; dots, individual data points (one-way ANOVA with Dunnett’s post hoc test compared to mock, p-value is represented; n.s.: no statistical significance).

The use of a GMI1000-derivative strain expressing green fluorescent protein (GFP) showed the presence of fluorescent bacteria in the meristematic/elongation zone following inoculation, which became particularly abundant at 48 and 72 hours post-inoculation (hpi), correlating with the inhibition of root growth (Figure S1K). It is important to note that, despite the fact that the *hrpG* mutant strain does not cause any significant alterations in Arabidopsis roots, it rapidly replicated in both root and shoot tissues after inoculation (Figures 1L and 1M), as usually observed in this *in vitro* experimental system (Yu et al, 2023), suggesting that the observed developmental alterations are not caused by the presence of bacterial cells multiplying in root tissues, but rather caused by specific T3SS-dependent activities.

### *R. solanacearum* affects root developmental programs

The development of the Arabidopsis root has been extensively studied, and multiple tools and experimental approaches are currently available to investigate root developmental events. Given that we have observed morphological changes in Arabidopsis roots inoculated with *R. solanacearum*, we decided to capitalize on one of the available toolboxes, the SAND lines (Marquès-Bueno *et al*., 2016), which allow monitoring the expression of marker genes real-time at cellular resolution, to gain insight into the physiological effects of *R. solanacearum* infection. In order to detect changes in a robust and unequivocal manner at early time points after inoculation, we used a relatively higher inoculation dose (10^7^ cfu/ml; (Lu *et al*., 2018)). In these conditions, Arabidopsis roots inoculated with GMI1000 displayed shorter roots (Figure S1A and S1B), enhanced root hair formation (Figure S1C and S1D), and earlier xylem differentiation (Figure S1E and S1F) compared to mock- or *hrpG*-inoculated roots, but also reduced meristem size (Figure S1G and S1H). In these experimental conditions, morphological alterations were barely detectable at 48 hours post-inoculation (hpi), while they became obvious after 54 hpi.

In order to monitor the expression of regulators of root development in a spatial and temporal manner, we combined staining with PI or FM4-64 and fluorescence microscopy upon *R. solanacearum* inoculation in different Arabidopsis lines expressing fluorescent reporters. WOX5 (WUSCHEL-RELATED HOMEOBOX 5) is a homeodomain transcription factor, and its expression in the QC is required for maintaining stem cell niche identity, contributing to the development of the root meristem (Sarkar *et al*., 2007). We did not detect alterations of the intensity or the spatial expression pattern of GFP driven by the *WOX5* promoter (*pWOX5::GFP*; (Blilou *et al*., 2005)), used as *WOX5* expression marker, at 1 or 2 days post-inoculation (dpi) (Figure 2A and 2D). However, seedlings inoculated with *R. solanacearum* GMI1000 WT showed a dramatic reduction in GFP fluorescence at 54 hpi compared with mock- or GMI1000 *hrpG*-inoculated roots (Figure 2A and 2D), indicating that *R. solanacearum* infection leads to a reduction of *WOX5* expression at the QC. SCARECROW (SCR) has been reported as an upstream regulator of *WOX5*, and it is required to specify QC identity and to maintain stem cells (Sabatini *et al*., 2003). SCR also contributes to the regulation of xylem patterning, participating in a regulatory pathway involving bidirectional cell signaling together with microRNAs 165 and 166 (miR165/6) and the transcription factor SHORT ROOT (SHR) (Carlsbecker *et al*., 2010). SCR maintains stem cell activities via directly repressing cytokinin-dependent response transcription factor ARABIDOPSIS RESPONSE REGULATOR 1 (ARR1) in the QC, which in turn controls auxin production, thus enabling stem cell division (Moubayidin *et al*., 2013). An Arabidopsis transgenic line expressing yellow fluorescent protein (YFP) under the *SCR* promoter (*pSCR::YFP*; (Marquès-Bueno *et al*., 2016)) shows YFP fluorescence at the endodermis and QC (Figure 2B and 2E). As observed for *pWOX5::GFP* (Figure 2A), there were no differences in the intensity or the spatial expression pattern of *pSCR::YFP* at 1 dpi to 2 dpi (Figure 2B and 2E). However, at 54 hpi, seedlings inoculated with *R. solanacearum* GMI1000 WT showed weaker fluorescence in the meristematic zone compared with mock- or GMI1000 *hrpG*-inoculated roots (Figure 2A). Altogether, these data show that *R. solanacearum* infection reduces the expression of *WOX5* and *SCR* at the root meristem, which may underlie the observed deficiencies in root meristem formation and primary root length upon *R. solanacearum* inoculation.

**Figure 2.**
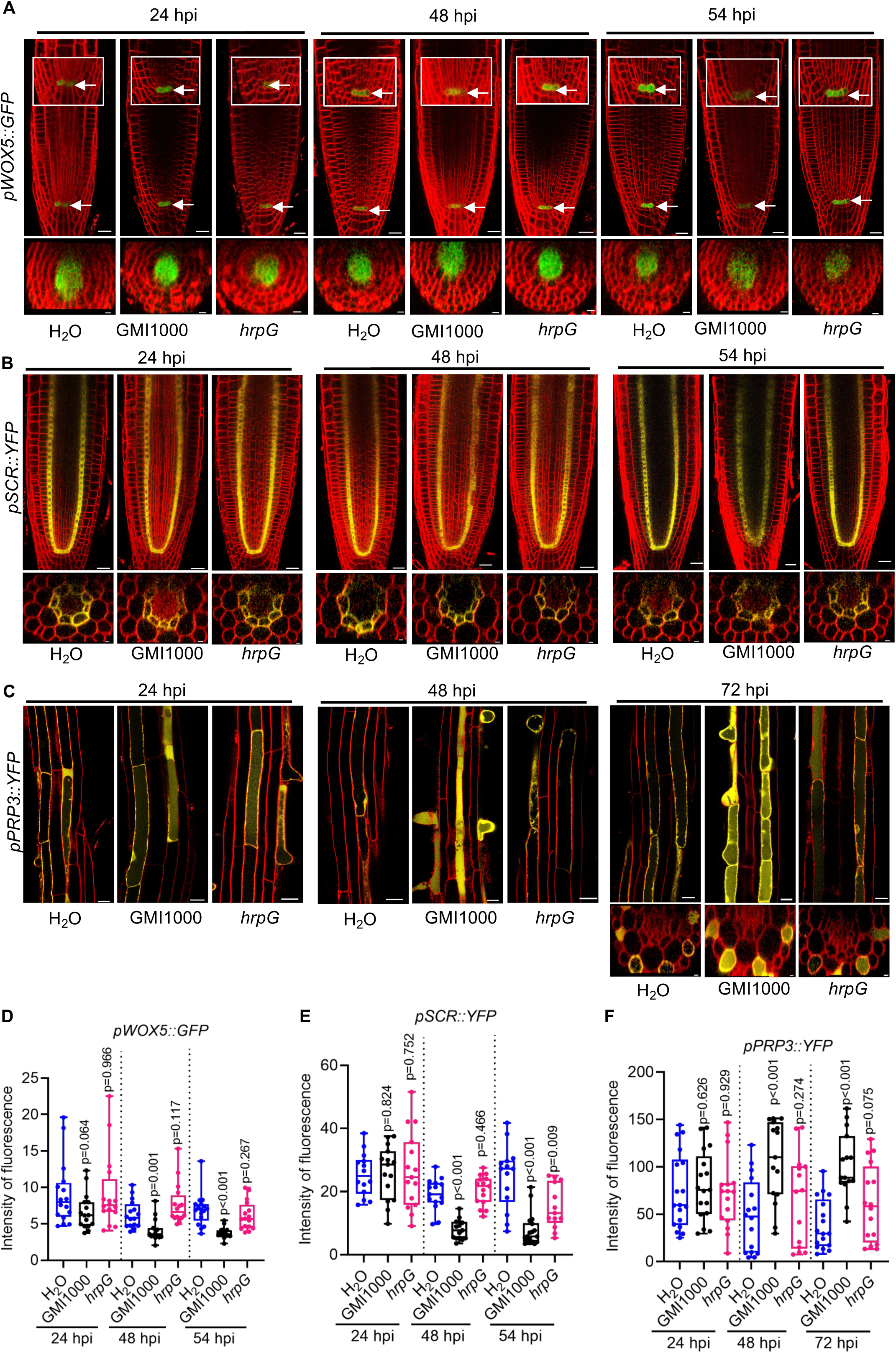
Inoculation with *Ralstonia solanacearum* alters root developmental programs. (A) *R. solanacearum* GMI1000 reduces the expression of *pWOX5::GFP* in the Arabidopsis root quiescent center (QC). Arrows represent QC. (B) *R. solanacearum* GMI1000 reduces the expression of *pSCR::YFP* in the Arabidopsis root meristem. (C) *R. solanacearum* GMI1000 enhances the expression of *pPRP3::YFP* in the Arabidopsis root maturation zone. (D-F) Quantification of fluorescence of *pWOX5::GFP*, *pSCR::YFP*, and *pPRP3::YFP* in (A-C), respectively. In (A-C), 7-d-old *pWOX5::GFP*, *pSCR::YFP*, and *pPRP3::YFP* Arabidopsis marker lines grown on MS^-^ plates were inoculated with different *R. solanacearum* strains (1×10^7^ cfu/ml), respectively. Seedlings were stained with FM4-64 (red), while the fluorescence was observed at indicated time point post inoculation (hpi, hours post-inoculation). Scale bars represent 20 μm for longitudinal and 5 μm for cross sections. At least 10 seedlings were used in three different experimental replicates with similar results. In all figure panels, GMI1000 means *R. solanacearum* GMI1000 wild-type strain; *hrpG* means *R. solanacearum* GMI1000 *hrpG* mutant strain; H_2_O means mock (in which water has been used instead of the bacterial inoculum). In (D-F), data pooled from three repeats are represented as boxes and whiskers, center line, median; box limits, lowest and highest data point; dots, individual data points (one-way ANOVA with Dunnett’s post hoc test compared to mock, p-value is represented; n.s.: no statistical significance).

We have shown that *R. solanacearum* infection promotes the formation of root hairs at the root tip. Root hairs are modified root epidermal cells that fortify the total absorptive surface of the root system and take part in water and nutrient uptake (Peterson & Farquhar, 1996). The Arabidopsis PROLINE-RICH PROTEIN 3 (PRP3) is a structural cell wall protein regulated by cell-type-specific developmental pathways involved in root hair formation (Bernhardt & Tierney, 2000). The spatiotemporal expression of *PRP3* can be monitored using Arabidopsis transgenic seedlings expressing the *pPRP3::YFP*, which show fluorescence in differentiating epidermal cells (Marquès-Bueno *et al*., 2016). Interestingly, seedlings inoculated with *R. solanacearum* GMI1000 WT showed stronger *pPRP3::YFP* fluorescence in epidermal cells of the maturation zone 2-3 dpi, compared with mock- and GMI1000 *hrpG*-inoculated seedlings, correlating with the promotion of root hair formation (Figure 2C and 2F). This indicates that *R. solanacearum* invasion promotes specific cell differentiation pathways that may lead to the observed alterations in root development.

Of note, the observed effects of *R. solanacearum* infection on the expression of *WOX5*, *SCR*, and *PRP3* are most likely specific, since other genes involved in root development, such as *MYB46* and *PIN4*, do not display alterations in their expression pattern or intensity (Figure S2A-S2F).

### *R. solanacearum* affects auxin accumulation and auxin-dependent signaling

Some of the morphological and molecular changes observed upon *R. solanacearum* inoculation in our previous experiments might be regulated by auxin, since this hormone regulates the expression of *WOX5* (Ding & Friml, 2010), root hair formation (Pitts *et al*., 1998; Grebe *et al*., 2002), and xylem differentiation (Fàbregas *et al*., 2015). Given the importance of auxin for root development and the fact that *R. solanacearum* infection induces auxin-dependent gene expression (Zuluaga *et al*., 2015; French *et al*., 2018; Zhao *et al*., 2019), we measured auxin accumulation in roots undergoing a *R. solanacearum* infection. As shown in Figure 3A, roots inoculated with GMI1000 accumulated more auxin than those treated with water or inoculated with the *hrpG* mutant strain, indicating that an active *R. solanacearum* infection induces auxin accumulation in roots.

**Figure 3.**
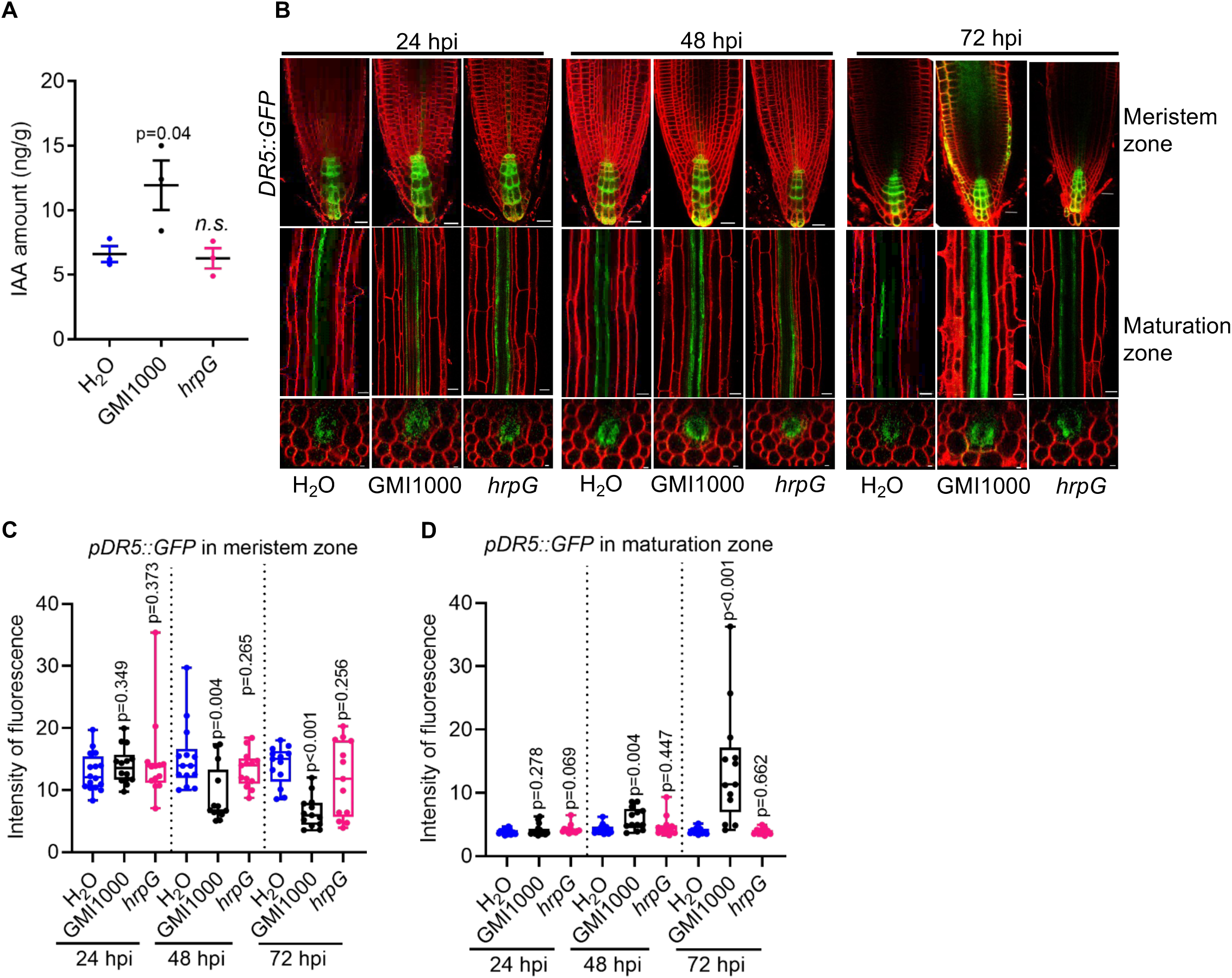
Inoculation with *Ralstonia solanacearum* alters auxin content and signaling in Arabidopsis. (A) Free Indole-3-acetic acid (IAA) levels in wild-type Col-0 roots inoculated with different strains. The inoculated root tissues (approx. 15 mg for each sample) were collected at 3 days post-inoculation. Data are represented as means ± SEM (n=3; one-way ANOVA with Dunnett’s post hoc test compared to mock, p-value is represented; n.s.: no statistical significance). (B) *R. solanacearum* GMI1000 enhances the expression of *DR5::GFP* in the root maturation zone and alters its spatial expression pattern. (C and D) Quantification of fluorescence of *DR5::GFP* in meristem zone and maturation zone in (B). In (A and B), 7-d-old Col-0, *DR5::GFP* Arabidopsis seedlings grown on MS plates were inoculated with different *R. solanacearum* strains (1×10 cfu/ml), respectively. In (B), seedlings were stained with FM4-64 (red), and the GFP fluorescence was observed at the indicated time point post inoculation (hpi, hours post-inoculation). Scale bars represent 20 μm for longitudinal in and 5 μm for cross sections. At least 10 seedlings were used in three different experimental replicates with similar results. In (C and D), data pooled from three repeats are represented as boxes and whiskers, center line, median; box limits, lowest and highest data point; dots, individual data points (one-way ANOVA with Dunnett’s post hoc test compared to mock, p-value is represented; n.s.: no statistical significance). In all figure panels, GMI1000 means *R. solanacearum* GMI1000 wild-type strain; *hrpG* means *R. solanacearum* GMI1000 *hrpG* mutant strain; H_2_O means mock (in which water has been used instead of the bacterial inoculum).

In order to investigate auxin responses in different root cell types during *R. solanacearum* infection, we inoculated roots of Arabidopsis transgenic plants expressing *DR5::GFP* (Friml *et al*., 2003). DR5 is a highly active synthetic auxin response element, which provides a powerful reporting system to study auxin-responsive transcription (Ulmasov *et al*., 1997). At early time-points upon inoculation (1 to 2 dpi), GFP fluorescence was detected toward the root tip in the root meristem and at the distal end of the vasculature, with no obvious differences between roots inoculated with *R. solanacearum* and mock-treated roots (Figure 3B), although signal quantification showed small but significant differences in roots inoculated with GMI1000 (Figure 3C and 3D). However, at 3 dpi, we observed stronger fluorescence in the maturation zone of roots inoculated with GMI1000 compared to mock- or *hrpG*-inoculated roots (Figure 3B-D). Interestingly, we also observed fluorescence in the meristem epidermal cells of roots inoculated with GMI1000 (Figure 3B and 3C), indicating that an active *R. solanacearum* infection not only enhances auxin accumulation and auxin responses at the root tip and in the vasculature, but also induces auxin responses in the epidermis.

### Auxin biosynthesis is required for the *R. solanacearum*-induced root hair formation, but not earlier xylem differentiation

To determine the requirement of auxin for the observed *R. solanacearum*-induced alterations of root development, we combined pharmacological and genetic approaches. L-Kynurenine (L-Kyn) is a tryptophan degradation product that blocks the activity of TRYPTOPHAN AMINOTRANSFERASE OF ARABIDOPSIS (TAA) aminotransferases, thus inhibiting auxin biosynthesis (He *et al*., 2011). Treatment with L-Kyn caused a reduction in root growth (Figure S2G), and, importantly, abolished the induction of *pPRP3::YFP* expression and the promotion of root hair formation upon *R. solanacearum* inoculation (Figure 4A and 4B). This is in agreement with the observation that the *R. solanacearum*-induced promotion of root hair formation is significantly compromised in Arabidopsis seedlings carrying a mutation in the auxin receptor TRANSPORT INHIBITOR RESPONSE 1 (TIR1) (Figure 4C and 4D; (Lu *et al*., 2018)). However, the earlier xylem differentiation induced by *R. solanacearum* infection was still observed in Arabidopsis WT seedlings treated with L-Kynurenine, despite the fact that root growth is inhibited to a similar length in the absence or presence of the bacteria, or Arabidopsis *tir1-1* mutant seedlings (Figure 4E). The treatment with L-Kyn in *tir1-1* mutant seedlings caused an important reduction of meristem size for all samples, without significant differences between mock, GMI1000 and the *hrpG* mutant (Figure 4F). These results indicate that auxin biosynthesis and perception is differentially required for distinct developmental alterations induced during *R. solanacearum* infection.

**Figure 4.**
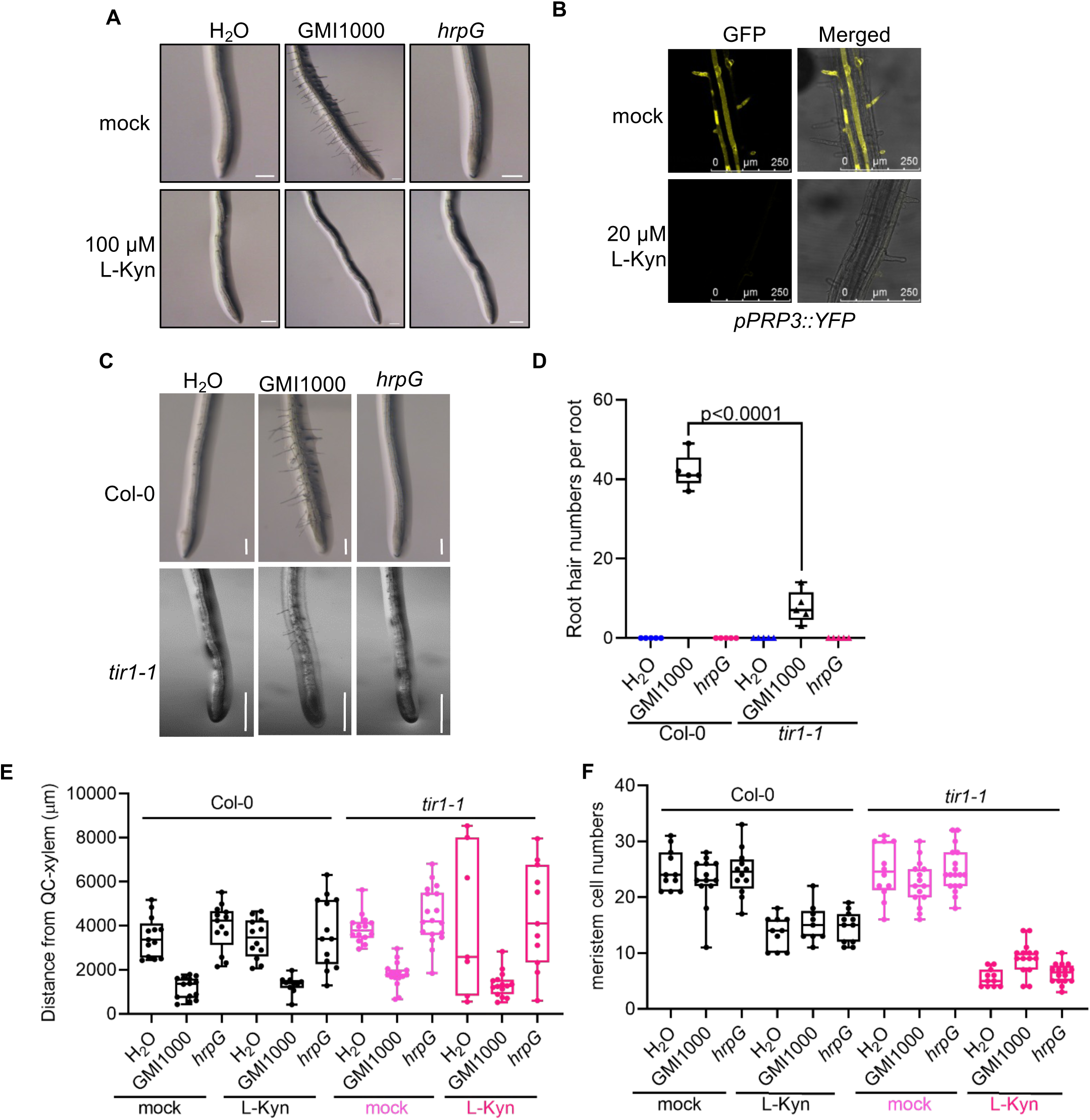
Auxin biosynthesis is required for *Ralstonia solanacearum*-induced root developmental changes. (A) Blocking auxin biosynthesis inhibits *R. solanacearum*-induced root hair formation. The auxin biosynthesis inhibitor L-Kynurenine (100 μM) was supplemented in the MS^-^ media and the pictures were taken 54 hours post-inoculation (hpi), scale bar=250 μm. (B) Blocking auxin biosynthesis inhibits the expression of *pPRP3::YFP* in Arabidopsis root. The auxin biosynthesis inhibitor L-Kynurenine (20 μM) was supplemented in the MS^-^ media and pictures were taken 48 hpi, scale bar=250 μm. (C and D) The *R. solanacearum-*mediated induction of root hair formation is diminished in Arabidopsis *tir1-1* mutant seedlings. In (C), pictures were taken at 54 hpi, scale bar=500 μm. (D) Quantification of root hairs in (C) (n=5 in each condition). Root hairs were counted in the 1.5-2 mm interval from the root tip in each root. (E) Blocking auxin biosynthesis does not affect *R. solanacearum*-induced earlier xylem differentiation in Arabidopsis wild-type or *tir1-1* mutant. The auxin biosynthesis inhibitor L-Kynurenine (L-Kyn, 20 μM) was supplemented in the MS media and the distance from QC to xylem was determined at 54 hpi. (F) Blocking auxin biosynthesis does not affect meristem cell numbers in Arabidopsis wild-type or *tir1-1* mutant. The auxin biosynthesis inhibitor L-Kynurenine (L-Kyn, 20 μM) was supplemented in the MS media and the meristem cell numbers were counted at 54 hpi. In (E and F), data are represented as pools of three repeats. In all experiments, 7 or 8-d-old Arabidopsis Col-0 or *tir1-1* seedlings on MS or ½ MS plates (specified in the methods section) were inoculated with different strains with a 1×10 cfu/ml inoculum. In all figure panels, GMI1000 means *R. solanacearum* GMI1000 wild-type strain; *hrpG* means *R. solanacearum* GMI1000 *hrpG* mutant strain; H_2_O means mock (in which water has been used instead of the bacterial inoculum). Each experiment was repeated two times (A, C, and D) and three times (B, E, F) with similar results. In (D-F), data are represented as boxes and whiskers, center line, median; box limits, lowest and highest data point; dots, individual data points. (t-test, p-value is represented).

### The developmental alterations induced by *R. solanacearum* do not require PEPR1/2 nor BAK1/BKK1

Plant invasion by bacterial pathogens is known to induce plant cellular damage. Even in the absence of obvious cell death, cellular damage may trigger the release of damage-associated molecular patterns (DAMPs), such as the small Pep peptides (Yamaguchi *et al*., 2006). In Arabidopsis, several Pep peptides can be perceived by the pattern-recognition receptors (PRRs) AtPEPR1 and AtPEPR2, leading to the activation of immune responses (Yamaguchi *et al*., 2006; Huffaker & Ryan, 2007; Krol *et al*., 2010; Yamaguchi *et al*., 2010). Moreover, treatment of Arabidopsis seedlings with Pep1 or Pep2 synthetic peptides has been shown to induce the formation of root hairs and an inhibition of primary root growth (Jing *et al*., 2019), mostly relying on perception by AtPEPR2 (Jing *et al*., 2019; Okada *et al*., 2021). Considering the overlap between *R. solanacearum*- and Pep1/2-induced root developmental alterations, we contemplated the possibility that cellular damage induced by *R. solanacearum* root invasion may underlie the observed root phenotypes. However, *R. solanacearum* still caused inhibition of root length, promotion of root hair formation, and earlier xylem differentiation in seedlings lacking both PEPR1 and PEPR2 ((Yamaguchi *et al*., 2010); Figure 5A-5D), indicating that root developmental alterations induced by *R. solanacearum* infection are not caused by the perception of endogenous DAMPs by these PRRs.

**Figure 5.**
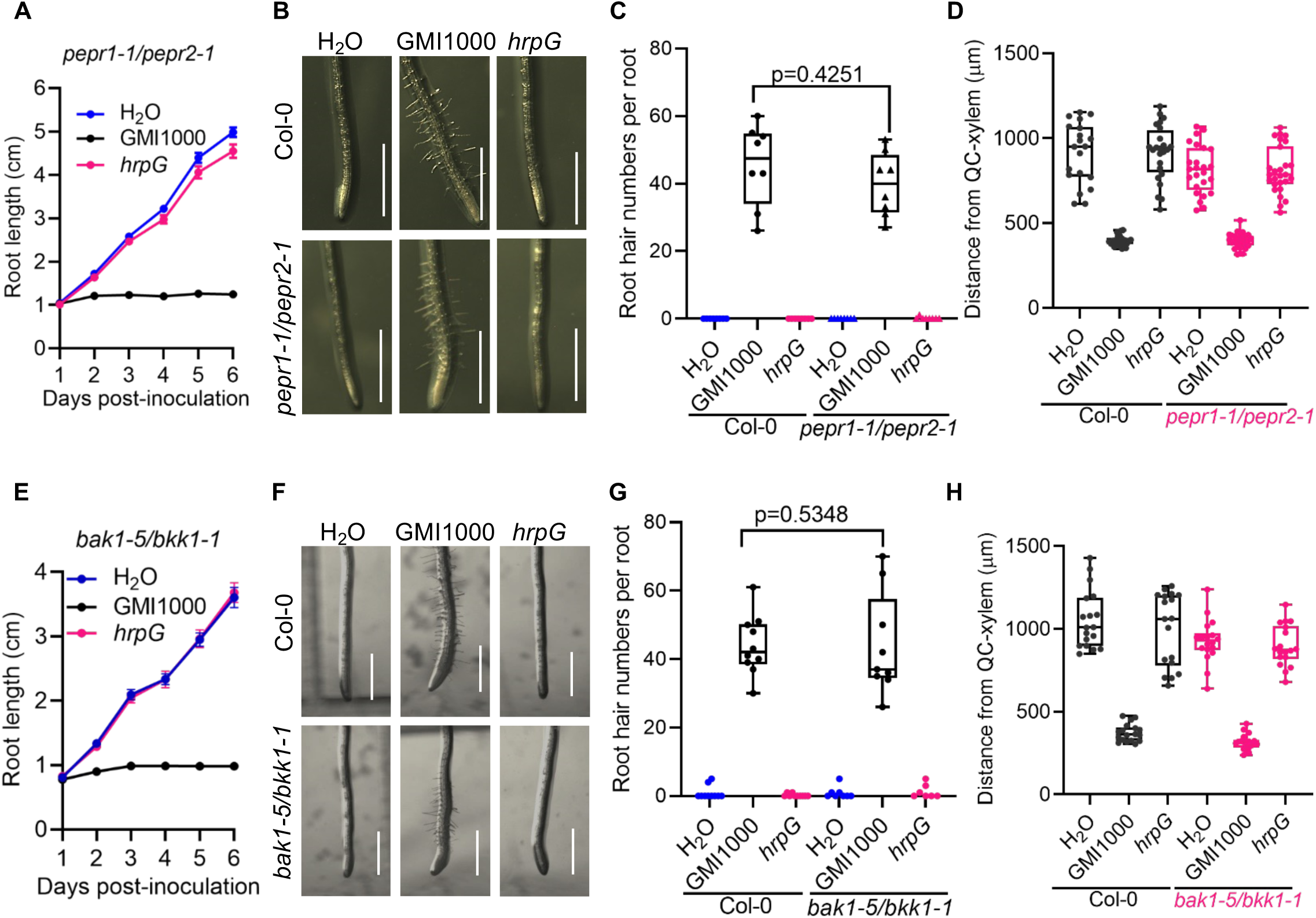
The developmental alterations induced by *Ralstonia solanacearum* do not require PEPR1/PEPR2 or BAK1/BKK1. (A) Time-course observations of primary root growth in *pepr1-1/pepr2-1* double mutant in different days post *R. solanacearum* inoculation. Data are represented as means ± SEM (n=22-24 in each condition). (B) *R. solanacearum* GMI1000 induces root hair formation in Arabidopsis. Pictures were taken at 54 hours post-inoculation (hpi), scale bar=1 mm. (C) Quantification of root hairs in (B) (n=8 in each condition). Root hairs were counted in the 1.5-2 mm interval from the root tip in each root. (D) Knock-out of *PEPR1* and *PEPR2* does not affect *R. solanacearum*-induced earlier xylem differentiation in Arabidopsis. The distance from QC to xylem was determined at 54 hpi, and data are represented as pools of three repeats. (E) Time-course observations of primary root growth in *bak1-5/bkk1-1* double mutant in different days post *R. solanacearum* inoculation. Data are represented as means ± SEM (n=19-29 in each condition). (F) *R. solanacearum* GMI1000 induces root hair formation in Arabidopsis. Pictures were taken at 54 hours post-inoculation (hpi), scale bar=1 mm. (G) Quantification of root hairs in (F, n=8-10 in each condition). Root hairs were counted in the 1.5-2 mm interval from the root tip in each root. (H) Knock-out of *BAK1* and *BKK1* does not affect *R. solanacearum*-induced earlier xylem differentiation in Arabidopsis. The distance from QC to xylem was determined at 54 hpi, and data are represented as pools of three repeats. In all experiments, 7-d-old Arabidopsis Col-0, *pepr1-1/pepr2-1*, or *bak1-5/bkk1-1* seedlings on MS-plates were inoculated with different strains with a 1×10^5^ cfu/ml inoculum. In all figure panels, GMI1000 means *R. solanacearum* GMI1000 wild-type strain; *hrpG* means *R. solanacearum* GMI1000 *hrpG* mutant strain; H_2_O means mock (in which water has been used instead of the bacterial inoculum). Each experiment was repeated three times with similar results. In (C,D, G and H), data are represented as boxes and whiskers, center line, median; box limits, lowest and highest data point; dots, individual data points. (t-test, p-value is represented).

Additionally, certain pathogen molecules can be perceived as pathogen-associated molecular patterns (PAMPs), which may induce a restriction in root growth (Parys *et al*., 2021). PAMPs, as DAMPs, are perceived by plasma membrane-localized PRRs (DeFalco & Zipfel, 2021). Most known PRRs require the co-receptor BAK1 and its paralog BKK1 for the activation of immune signaling upon elicitor perception (Zhou & Zhang, 2020). Interestingly, the developmental alterations observed upon *R. solanacearum* inoculation, including inhibition of root length, promotion of root hair formation, and earlier xylem still took place in a *bak1-5/bkk1-1* double mutant (Figure 5E-H), suggesting that root developmental alterations induced by *R. solanacearum* infection are not caused by the perception of bacterial PAMPs by BAK1/BKK1-dependent PRRs.

## DISCUSSION

Pathogen infection induces developmental alterations in host plants. In some cases, these alterations are considered disease symptoms caused by pathogen proliferation, which have detrimental effects on plant fitness, and may eventually cause the death of the infected organ or the whole plant. However, in other cases, developmental changes may be actively induced by the pathogen as part of its virulence activity, in order to facilitate its own proliferation and/or colonization of the host. In most cases, the causes and functions of these developmental alterations are to date unclear.

In this work, we found that infection by *R. solanacearum* alters root developmental programs and modifies root architecture, including the induction of root hair formation and premature xylem development near the root tip. A mutant bacterial strain lacking the major virulence regulator HrpG (which is required for the expression of the components of the T3SS, among other virulence factors) does not induce significant developmental alterations despite being able to reach population numbers similar to WT GMI1000 inside the plant in the experimental conditions used (Figure 1L-M), suggesting that these alterations are a product of bacterial virulence activities rather than collateral consequences of the infection. Moreover, treatment with synthetic PAMPs, such as the flagellin epitope flg22, does not induce similar changes in root architecture (Millet *et al*., 2010; Ranf *et al*., 2011; Beck *et al*., 2014). Finally, we observed similar *R. solanacearum*-induced developmental alterations in a *bak1-5/bkk1-1* mutant, which lacks responsiveness to most well-studied bacterial PAMPs (Ma *et al*., 2016), suggesting that they are not a consequence of the recognition of bacterial PAMPs during *R. solanacearum* infection. It was recently reported that endogenous Pep peptides, which act as DAMPs perceived by the receptors PEPR1 and PEPR2, induce root growth inhibition and root hair formation (Jing *et al*., 2019; Okada *et al*., 2021). Interestingly, *PEPR2* expression in certain root tissues is sufficient to trigger root growth inhibition and root hair formation in response to Pep1, but not to activate Pep1-triggered defence responses (Okada *et al*., 2021), uncoupling these developmental alterations from the induction of immunity. The observation that the root changes induced by *R. solanacearum* also take place in the absence of PEPR1 and PEPR2 (Figure 5) further supports the notion that these changes are a consequence of bacterial virulence activities rather than the production of cellular damage and the subsequent perception of endogenous DAMPs.

The observed modified development may provide an advantage for bacterial invasion and colonization of host plants. On one hand, the formation of additional root hairs may provide new entry points for these soil-borne bacteria; on the other, an earlier or enhanced proliferation of xylem vessels may increase the physical space for bacterial colonization and proliferation in the root vascular system, since xylem vessels are the main niche where *R. solanacearum* multiplies and the channel for this pathogen to gain access to the rest of the plant. Therefore, these alterations may benefit *R. solanacearum* at the population level.

Different plant pathogens have been shown to stimulate auxin synthesis in the plant they infect, or can synthesize auxin by themselves, what is believed to enhance susceptibility (reviewed in (Naseem *et al*., 2015)). The observation that treatment with the auxin biosynthesis inhibitor L-Kynurenine abolishes the *R. solanacearum*-induced *PRP3* overexpression and root hair development suggests that the increase in auxin observed during the infection is probably plant-derived. Of note, the earlier xylem differentiation is maintained in conditions of impaired auxin synthesis, which indicates that the developmental changes triggered by this pathogen are both auxin-dependent and -independent, and perhaps the result of multiple virulence activities exerted by different bacterial virulence factors.

In summary, our results indicate that invasion by the soil-borne pathogen *R. solanacearum* alters plant root development, and suggest that this bacterium has evolved virulence factors to manipulate this process through at least two independent strategies. It is worth noting that this study was performed using controlled experimental conditions in order to provide an accuracy and reproducibility that cannot be obtained in a natural field infection. Although the techniques used here to replicate the infection process are already non-invasive, further applied research will be required to determine the impact of all laboratory work in the more complex natural field conditions. Moreover, the future elucidation of the molecular mechanisms underpinning the potential co-option of plant development by the pathogen will not only shed new light on bacterial virulence strategies, but may also open new avenues for the generation of resistance to the devastating bacterial wilt disease.

## MATERIALS AND METHODS

### Plant materials and growth

The *Arabidopsis thaliana* materials used in this study are in ecotype Col-0, including Col-0 wild-type, *pWOX5::GFP* (Blilou et al., 2005), *DR5::GFP* (Friml *et al*., 2003), SAND lines (*pSCR::YFP*, *pPRP3::YFP*, *pMYB46::YFP*, and *pPIN4::YFP*; (Marquès-Bueno *et al*., 2016)), *tir1-1* (Ruegger *et al*., 1998), *pepr1-1/pepr2-1* (Yamaguchi *et al*., 2010), and *bak1-5/bkk1-1* (Roux *et al*., 2011).

Sterilized seeds were sown on MS^-^ plates (4.4 g L^-1^ Murashige and Skoog medium with vitamins, 0.5 g L^-1^ MES, 8 g L^-1^ bacto agar, pH 5.8) (Lu *et al*., 2018) and kept at 4°C for 2 days for stratification. Plates were then transferred to growth chambers (22°C, 16 h light/8 h dark, 100-150 mE m^-2^ s^-1^) for germination and vertical growth. Seedlings were grown for 7 days before bacterial inoculation. For chemical treatments, 5-d-old Arabidopsis seedlings were transferred to MS^-^ plates supplemented with the auxin biosynthesis inhibitor L-Kynurenine (L-Kyn, 20 or 100 μM) for 3 more days before bacterial inoculation. Due to the poor germination and growth of *tir1-1* on MS^-^ plates (with no sucrose), the experiments using the *tir1-1* mutant (in Figure 4) were performed as following: sterilized seeds were sown on ½ MS plates for 5 days, and then transferred to MS^-^ plates for 3 more days before bacterial inoculation.

### Bacterial strains and growth

*Ralstonia solanacearum* strains, including the phylotype I reference strain GMI1000 wild-type (Salanoubat *et al*., 2002), GMI1000 *hrpG* mutant (Valls *et al*., 2006), and GFP-expressing GMI1000 (GMI1000-GFP, Cao *et al*., 2023) were used in this study, and sterilized H_2_O was used as mock inoculation. Bacterial strains were first streaked out on solid Bacto-agar and glucose (BG) medium (Plener *et al*., 2010) for two days and a single colony was inoculated in complete BG liquid medium at 28 °C. The overnight cell culture was harvested by centrifugation (5000 g, 10 minutes) and re-suspended in sterilized de-ionized H_2_O (ddH_2_O). The bacterial inoculum was adjusted with ddH_2_O to high dose (1×10^7^ cfu/ml) and low dose (1×10^5^ cfu/ml) for inoculation.

### Bacterial inoculation

Bacterial inoculation was performed as previously described (Yu *et al*., 2023). Seven or eight day-old Arabidopsis seedlings were inoculated with 5 μl of a bacterial suspension at the root tip (approx. 0.5-1 cm from the root tip) and the plates were kept for 10 min in a sterile flow hood to let inoculation spots dry. Plates with inoculated seedlings were then transferred to a controlled growth chamber (75% humidity, 12 h light, 130 mE m^-2^ s^-1^, 27 °C) for vertical growth. Inoculated seedlings were used for subsequent experiments at the indicated time points.

### Staining

Modified pseudo-Schiff propidium iodide (mPS-PI) staining was performed as previously described (Truernit *et al*., 2008). Briefly, whole seedlings were fixed in fixative buffer (50% methanol and 10% acetic acid) at 4°C for at least 12 hours. Next, tissues were rinsed with water and incubated in 1% periodic acid at room temperature for 40 minutes. The tissue was rinsed again with water and incubated in Schiff reagent with propidium iodide (PI; freshly prepared PI to a final concentration of 100 μg mL^-1^) for 1 to 2 hours or until plants were visibly stained. The samples were then transferred to chloral hydrate solution (4 g chloral hydrate, 1 ml glycerol, and 2 ml water). Finally, the samples were used straightaway or kept at 4°C for 1 week.

Sample preparation for the observation of the Casparian strip in Arabidopsis was performed according to (Naseer *et al*., 2012). Inoculated seedlings were transferred to small Petri dishes containing 240 mM HCl in 20% methanol and incubated on a 57°C heat block for 15 minutes. Then the solution was replaced with 7% NaOH, 7% hydroxylamine-HCl in 60% ethanol for 15 minutes at room temperature. Then seedlings were rehydrated for 5 minutes each in 40%, 20%, and 10% ethanol, and infiltrated for 15 minutes in 5% ethanol and 25% glycerol. Finally, seedlings were mounted in 50% glycerol on glass microscope slides.

The fluorescent marker lines were stained with FM4-64 for better visualization of the cells. To perform FM4-64 staining, seedlings were treated with 1 μM FM4-64 solution (Life Technologies, stock solution at 1 mM in H_2_O) for 10-15 minutes at room temperature. Stained roots were incubated in water for 20-40 minutes before microscopic observation.

### Phenotypic and microscopic observation

To measure the impact of *R. solanacearum* on Arabidopsis root elongation, photographs were taken with a Nikon D7000 camera over time, and the primary root length was determined using the ImageJ software. To record root hair formation, seedlings grown on MS^-^ plates were observed 54 hours post-inoculation (hpi) using a Leica M205 FCA fluorescence stereo microscope. To analyze PI-stained or FM4-64 stained roots, root tissues were observed using a Leica TCS SP8 point scanning confocal microscope or a Leica TCS SMD FLCS single molecule detection system. To determine the root meristem sizes and the distance from the QC (quiescent center) to vascular xylem elements, confocal images of PI-stained roots were analyzed using the ImageJ software. Root meristem size was assessed as the number of cortical cells between the QC and the first cell that was twice the length of the immediately preceding cell (Perilli & Sabatini, 2010), while the distance from QC to vascular xylem elements was measured based on the distance from QC to the first appearance of differentiated xylem, using ImageJ. To determine auto-fluorescence of Casparian strip, the prepared root tissues were observed using a Leica TCS SP8 point scanning confocal microscope with GFP fluorescence settings, while the distance from the QC to the first appearance of the Casparian strip was analyzed using ImageJ. To observe the effect of L-Kynurenine on *pPRP3::YFP* expression, 5-d-old *pPRP3::YFP* seedlings were transferred to MS^-^ plates supplemented with 20 μM L-Kyn and the YFP fluorescence was observed 48 hours after incubation using a Leica TCS SP8 point scanning confocal microscope. To observe bacterial progression in Arabidopsis roots, the GFP-expressing strain (GMI1000-GFP) was inoculated on Col-0 root tips as described above, and the inoculated roots were washed three times in clean water before GFP fluorescence determination. GFP fluorescence in root tips and the differentiation zone was observed using a Leica TCS SP8 point scanning confocal microscope with GFP settings. The settings for the laser scanning are as follows: Ex: 488 nm, Em: 500-550 nm for GFP; Ex: 514 nm, Em: 525-570 nm for YFP; Ex: 561 nm, Em: 630-680 nm for PI staining; Ex: 561 nm, Em:650-695 nm for FM4-64 staining. The determination of the fluorescence intensity was conducted with ImageJ.

### Auxin measurement

Three days post-inoculation, root tissues from approximately 45 seedlings were collected in 1.5 mL Eppendorf tubes and immediately frozen in liquid nitrogen. Samples were then ground into powder using a Qiagen tissue lyser at 30 Hz for 1min. A total of 0.5 mL 70% methanol containing 1 ng [^13^C_6_]-IAA (CLM-1896-0.01, Cambridge Isotope Laboratories, USA), as an internal standard, was added into the tube, which was vortexed at 1000 rpm at 10 °C for 1 h and then centrifuged at 20000 g at 20 °C for 10 min. 0.3 mL supernatant was taken and diluted 2 times with H_2_O in a new HPLC vial. 50 µL solution was injected into the Acquity Ultra High Performance Liquid Chromatography/Mass Spectroscopy (UPLC/MS, Waters, USA) on a BEH C18 column at the Proteomics and Metabolomics Core Facility of the Shanghai Center for Plant Stress Biology. IAA contents were determined by comparing IAA peak areas with those of [^13^C_6_]-IAA (Wang *et al*., 2020). IAA contents (ng IAA per g of fresh root tissues) were calculated for statistical analysis.

### Data analysis

The generation of graphs and statistical analyses were performed using the Graphpad prism software.

## ACKNOWLEDGEMENTS

We thank Drs. Cyril Zipfel, Lin Xu, Haibin Lu, Meixiang Zhang, and Shingo Nagawa for sharing biological materials, Xinyu Jian and Fangyuan Wu for technical and administrative assistance during this work, and all the members of the Macho and Lozano-Duran laboratories for helpful discussions. We thank the PSC Cell Biology core facility for assistance with microscopy assays. This work was supported by the Strategic Priority Research Program of the Chinese Academy of Sciences (grant XDB27040204), the National Natural Science Foundation of China (NSFC; grants 32170289 to APM, and 32270284 to GY), the Chinese 1000 Talents Program, and the Shanghai Center for Plant Stress Biology (Chinese Academy of Sciences). The authors have no conflict of interest to declare.

## AUTHOR CONTRIBUTIONS

RLD and APM planned and designed the research. LZ, GY, HX, and ML performed experiments and analysed data. RLD and APM wrote the manuscript.

---

**Figure S1.**
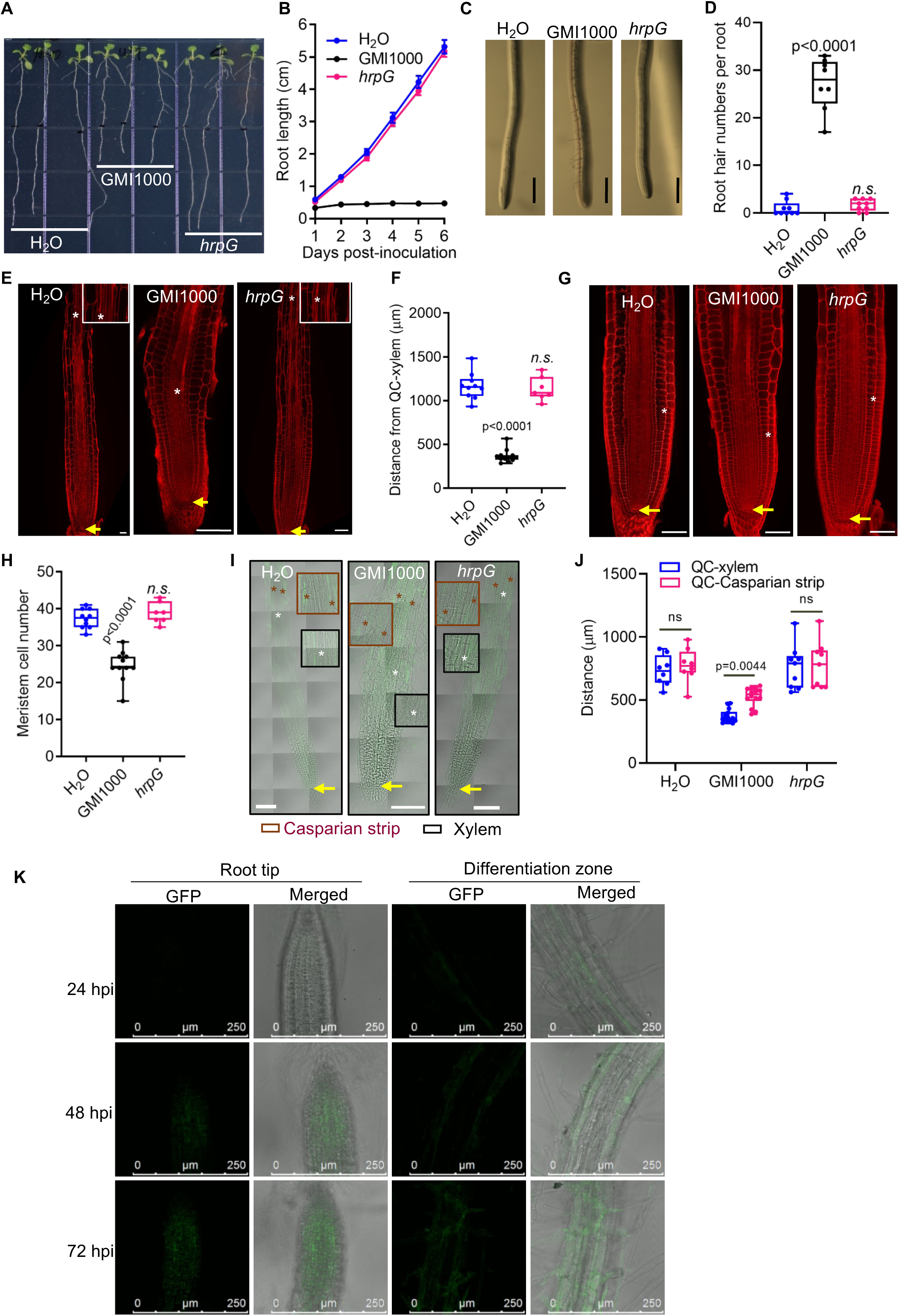
Inoculation with *Ralstonia solanacearum* using a high dose causes Arabidopsis root development alterations. (A) *R. solanacearum* GMI1000 inhibits primary root growth in Arabidopsis. Pictures were taken at 6 days post-inoculation (dpi). (B) Time-course observations of primary root growth in different days post-inoculation. Data are represented as means ± SEM (n=8 in each condition). (C) *R. solanacearum* GMI1000 induces root hair formation in Arabidopsis. Pictures were taken 54 hours post-inoculation (hpi), scale bar=500 μm. (D) Quantification of root hairs in (C) (n=8 in each condition). Root hairs were counted in the 1.5-2 mm interval from the root tip in each root. (E and F) *R. solanacearum* GMI1000 causes earlier xylem differentiation in Arabidopsis. In (E) the inoculated seedlings were used for propidium iodide (PI) staining at 3 days post-inoculation (dpi). Yellow arrows and white asterisks indicate the QC and the first appearance of xylem, respectively, scale bar = 100 μm. In (F), the distance from the QC to the xylem was measured from (E), and data represents one of the replicates. (G and H) *R. solanacearum* GMI1000 causes a reduction of root meristem cell numbers in Arabidopsis. In (G) the inoculated seedlings were used for PI staining at 3 dpi. Yellow arrows and white asterisks indicate the QC and the first cortical cell that was twice the length of the preceding cell, respectively, scale bar=100 μm. In (H), meristem cells were counted and data represents one of the replicates. (I) *R. solanacearum* GMI1000 causes earlier xylem and Casparian strip differentiation in Arabidopsis, yellow arrows, white asterisks, and red asterisks indicate the quiescent center (QC) and the first appearance of xylem and Casparian strip, respectively, scale bar = 100 μm. (J) The distance from the QC to the xylem or the first appearance of Casparian strip in (I), and data are represented as pools of three repeats (H_2_O: n=8; GMI1000: n=14; *hrpG*: n= 9). (K) Observations of GFP fluorescence in Arabidopsis root at different time points post-*Ralstonia* inoculation. 7-old Arabidopsis Col-0 seedlings on MS^-^ plates were inoculated with *R. solanacearum* GMI1000 expressing GFP (GMI1000-GFP) (CFU=1×10^7^ cfu/ml), and the GFP fluorescence in root tip and differentiation zone was observed at different time points, scale bar = 250 μm. In all experiments (A-H, and K), 7-d-old Arabidopsis Col-0 seedlings on MS^-^ plates were inoculated with different *R. solanacearum* strains with a dose (1×10^7^ cfu/ml) higher to that used in Figure 1 (1×10^5^ cfu/ml), while in (I-J) the experimental conditions are the same as in main Figure 1J-1K. In all figure panels, GMI1000 means *R. solanacearum* GMI1000 wild-type strain; *hrpG* means *R. solanacearum* GMI1000 *hrpG* mutant strain; H_2_O means mock (in which water has been used instead of the bacterial inoculum). Each experiment was repeated two (A-H) or three (I-K) times with similar results. In (D, F, and H), data are represented as boxes and whiskers, center line, median; box limits, lowest and highest data point; dots, individual data points (one-way ANOVA with Dunnett’s post hoc test compared to mock, p-value is represented, n.s. corresponds to no statistical significance).

**Figure S2.**
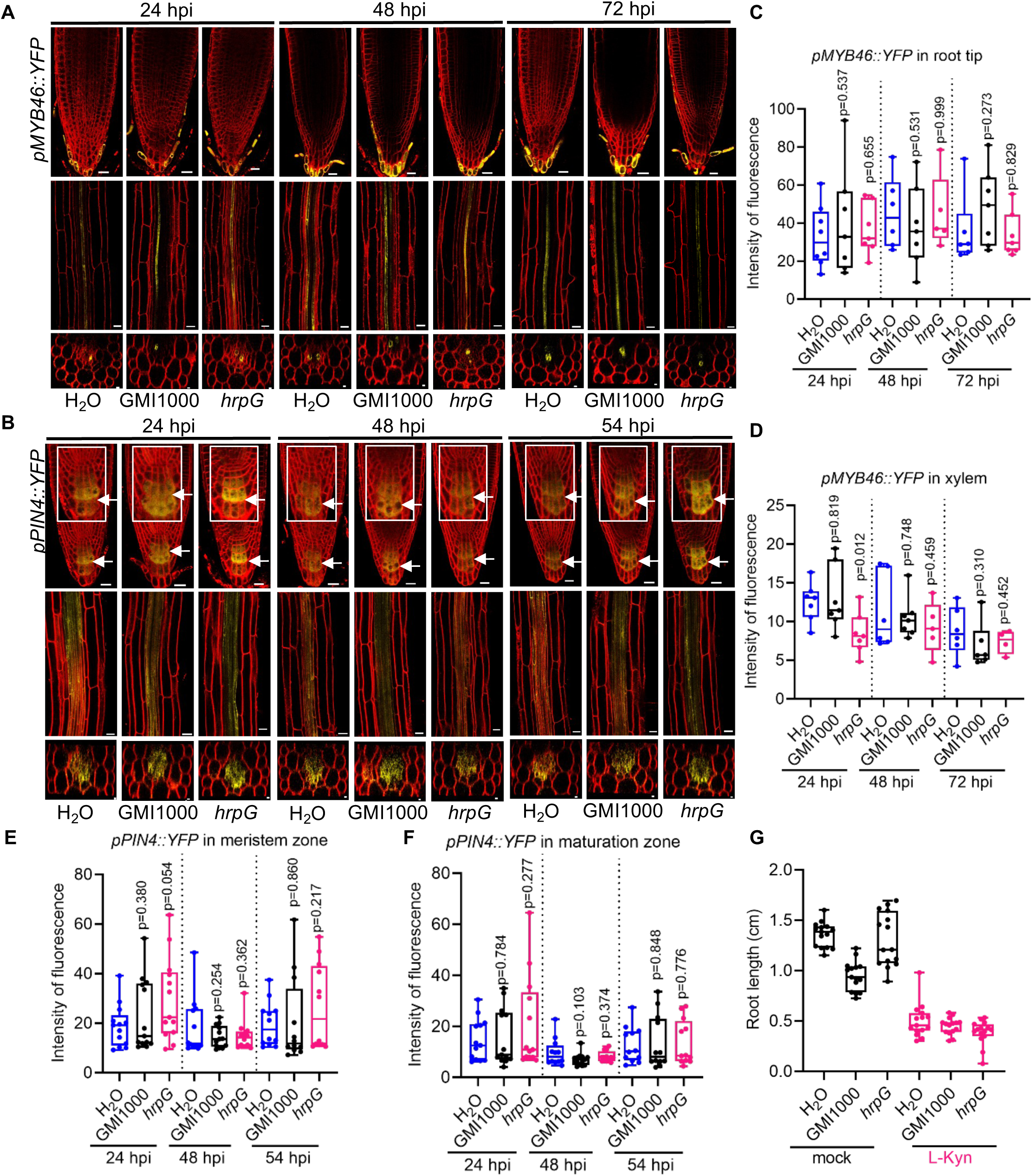
Inoculation with *Ralstonia solanacearum* does not affect *MYB46* and *PIN4* expression in the Arabidopsis root. (A) and (B) *R. solanacearum* GMI1000 does not affect the expression of *pMYB46::YFP* or *pPIN4::YFP* in the Arabidopsis root. (C and D) Quantification of *pMYB46::YFP* fluorescence in root tip and xylem tissue in (A). (E and F) Quantification of *pPIN4::YFP* fluorescence in meristem zone and maturation zone in (B). In (A and B), 7-d-old *pMYB46::YFP* and *pPIN4::YFP* Arabidopsis marker lines grown on MS^-^ plates were inoculated with different *R. solanacearum* strains (1×10^7^ cfu/ml), respectively. Seedlings were stained with FM4-64 (red), and the YFP fluorescence was observed at the indicated time point post inoculation (hpi, hours post-inoculation). Scale bars represent 20 μm for longitudinal and 5 μm for cross sections. 8 to 10 seedlings were used in two different experimental replicates with similar results. (G) Quantification of primary root length of Arabidopsis Col-0 in Figure 4E. The auxin biosynthesis inhibitor L-Kynurenine (L-Kyn, 20 μM) was supplemented in the MS^-^ media and the primary root length was determined at 54 hpi, and data are represented as pools of three repeats (n=15). In all figure panels, GMI1000 means *R. solanacearum* GMI1000 wild-type strain; *hrpG* means *R. solanacearum* GMI1000 *hrpG* mutant strain; H_2_O means mock (in which water has been used instead of the bacterial inoculum). In (C-G), data are represented as boxes and whiskers, center line, median; box limits, lowest and highest data point; dots, individual data points (one-way ANOVA with Dunnett’s post hoc test compared to mock, p-value is represented, n.s. corresponds to no statistical significance).

